# Cell-free and *in vivo* characterization of Lux, Las, and Rpa quorum activation systems in *E. coli*

**DOI:** 10.1101/159988

**Authors:** Andrew D. Halleran, Richard M. Murray

## Abstract

Synthetic biologists have turned towards quorum systems as a path for building sophisticated microbial consortia that exhibit group decision making. Currently, however, even the most complex consortium circuits rely on only one or two quorum sensing systems, greatly restricting the available design space. High-throughput characterization of available quorum sensing systems is useful for finding compatible sets of systems that are suitable for a defined circuit architecture. Recently, cell–free systems have gained popularity as a test-bed for rapid prototyping of genetic circuitry.

We take advantage of the transcription-translation cell-free system to characterize three commonly used Lux-type quorum activators, Lux, Las, and Rpa. We then compare the cell-free characterization to results obtained *in vivo.* We find significant genetic crosstalk in both the Las and Rpa systems and substantial signal crosstalk in Lux activation. We show that cell-free characterization predicts crosstalk observed *in vivo*.

## Main text

Multicellular systems achieve extraordinary complexity through specialization and communication between cell types to orchestrate high-level behaviors. Over the last 15 years, synthetic biologists have begun to utilize bacterial communication systems for orchestrating behavior at the group level instead of individual cells. Quorum sensing (QS) systems are simple bacterial circuits that allow coordination of group behaviors within local populations of bacteria. ^1, 2^ Early synthetic circuits exploited the Lux system to build a population control device that artificially capped a growing *E. coli* population.^3^ Building off this work, a more complex “predator-prey” community was built where two distinct cell types each expressed one of the Las or Lux quorum systems.^4^ This synthetic community could produce extinction, coexistence, and oscillatory behaviors. Recent efforts have investigated how circuit topologies within microbial communities influence the stability of an example emergent property, oscillations.^5^

Expanding this class of circuits to more complex architectures currently remains challenging to engineer due to significant crosstalk between QS systems. Commonly-used Lux-type QS systems use a transcription factor sensitive to a small hydrophobic molecule, acyl homoserine lactone (AHL), produced by an AHL synthase, that can then activate a target promoter containing a lux-box response element.^6^ Lux-type QS systems share significant similarity at the transcription factor, promoter, and ligand level.^6^ This has hindered development of synthetic circuits that exploit simultaneous expression of more than one or two QS systems.

Recent work has attempted to both characterize and reduce this crosstalk. Hasty and colleagues examined four Lux-type quorum activators, Lux, Las, Rpa, and Tra, and characterized observed crosstalk patterns *in vivo.*^7^ Work by Haseloff and colleagues engineered the canonical pLux and pLas promoters to reduce crosstalk, allowing for complex pattern-generating circuits.^8^

Characterizing these combinatorial design spaces becomes increasingly challenging as the number of elements (transcription factor, ligand, promoter, synthase) and systems (Lux, Las, …) of interest increases. The cell-free expression system TX-TL has been used successfully to quickly prototype and iterate complex circuits that would be difficult to build and test *in vivo*, and previous work has shown that Las activation in a cell-free system is predictive of *in vivo* performance.^9–13^ To increase the potential characterization space, we tested whether one could characterize quorum systems using TX-TL as a rapid-prototyping platform.

We chose to test three commonly used Lux-type quorum transcription factors, LuxR, LasR, RpaR, their cognate promoters (pLux, pLas, pRpa) and cognate AHLs (3OC6, 3OC12, p-coumaroyl) for crosstalk in TX-TL. To investigate both signal crosstalk, which is mediated by transcription factors activated by non-cognate AHLs, and genetic crosstalk, which is caused by active transcription factor — AHL complexes targeting non-cognate promoters, we tested all combinations of transcription factor, AHL, and promoter (27 total) at 8 different concentrations of AHL (216 combinations). All transcription factors were cloned into a standard plasmid cassette using a constitutive promoter (BBa_J23106), and a strong ribosome binding site (BBa_K1114107) onto a high-copy backbone.^14, 15^ TX-TL reactions were prepared as previously described and conducted in a plate reader to collect time course fluorescence measurements. An example on-target activation of RpaR, p-coumaroyl, and pRpa-GFP is shown in Figure 1A (all traces can be found in supporting information figures S1-S27).

**Figure 1:**
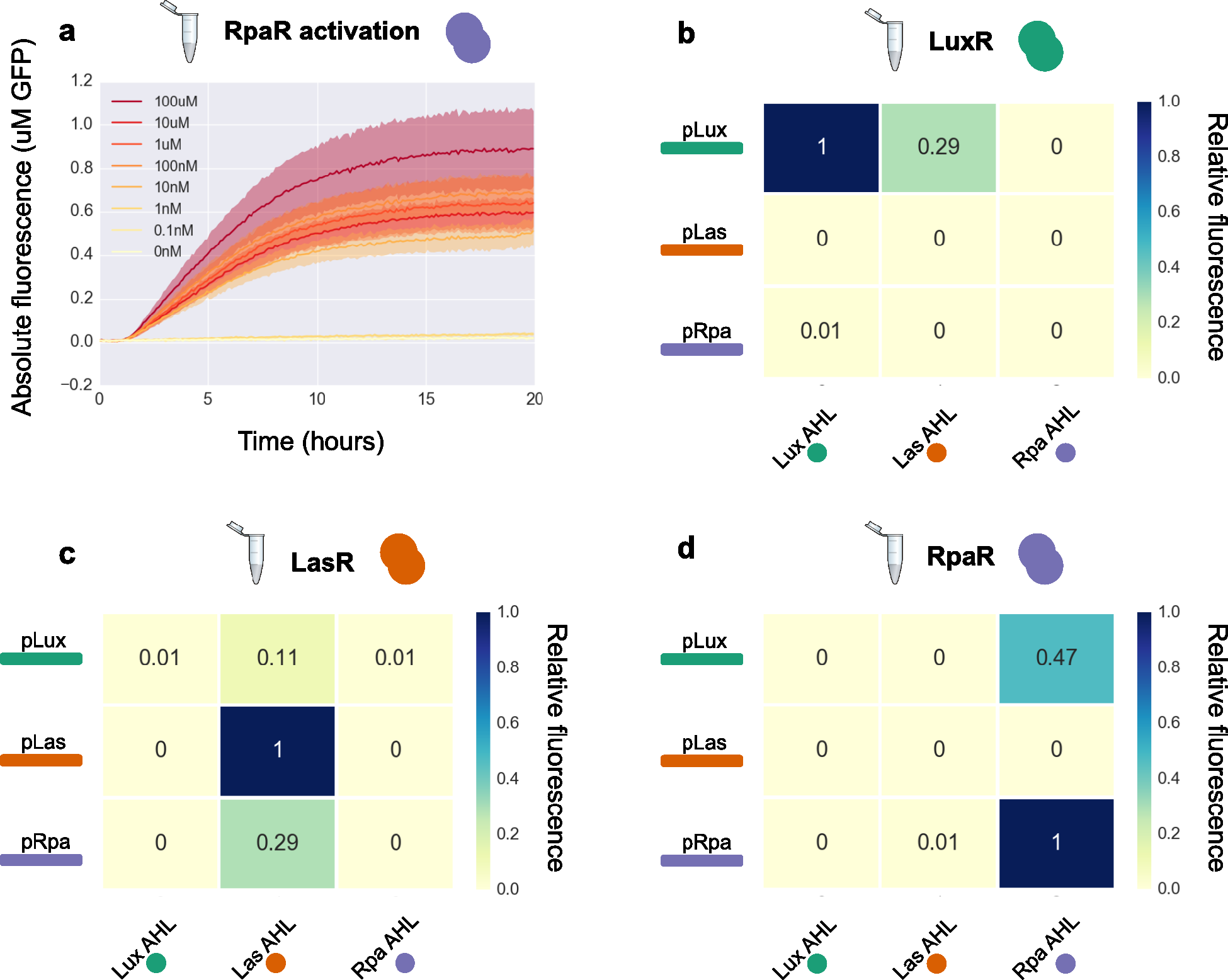
TX-TL crosstalk mapping. (A) Example time trace from RpaR (2nM DNA) + pRpa-GFP (4nM DNA) across a range of p-coumaroyl concentrations. Solid lines represent the mean of four replicates, shading represents +/- one standard deviation. (B) Crosstalk heatmap for LuxR (2nM LuxR), with all three reporters (4nM each), and AHLs (1uM). Relative fluorescence is calculated for each combination as the average of the 95^th^ percentile fluorescence value across four replicates divided by the maximum observed across all nine combinations. (C) Crosstalk heatmap for LasR. (D) Crosstalk heatmap for RpaR.

To characterize both genetic and signal crosstalk we tested all 27 combinations of transcription factor, AHL, and promoter in TX-TL. Traces from 1uM AHL concentration were used to calculate crosstalk heatmaps. LuxR is activated by both Lux AHL (3OC6) and Las AHL (3OC12) (Figure 1B). However, regardless of the activating AHL, LuxR retains genetic specificity to the pLux promoter, showing no significant crosstalk with pLas or pRpa. In stark contrast with LuxR, LasR shows substantial genetic promiscuity, targeting both the pLux and pRpa promoters (Figure 1C). LasR has no discernible signal-level crosstalk, only showing activation of target promoters when induced with its native ligand. RpaR, like LasR, is only activated by its cognate ligand (p-coumaroyl), and displays a distinct genetic crosstalk pattern where it strongly activates both pRpa and pLux, but does not activate pLas (Figure 1D).

To determine how predictive our cell-free characterizations were of *in vivo* circuit results, we double-transformed all nine combinations of activator and reporter plasmids into *E. coli.* Despite accurately predicting observed signal crosstalk patterns for all three systems, cell-free experiments underestimated the amount of genetic crosstalk for both LuxR and LasR (Figure 2A-B). In the case of LuxR, TX-TL predicted that there would be no detectable genetic crosstalk with the other two promoters, but *in vivo* results showed that LuxR + Lux AHL readily activated both pLux and pRpa, while failing to activate pLas. LasR *in vivo* results also showed increased amounts of genetic crosstalk compared to TX-TL, but the same pattern of high genetic crosstalk and low signal crosstalk predicted by cell-free experiments was confirmed *in vivo.* RpaR shows an extremely similar mapping, both qualitatively and quantitatively, from TX-TL to *in vivo* results (Figure 2C). Both the lack of signal crosstalk, and the genetic crosstalk with pLux, was demonstrated in cell-free experiments.

**Figure 2:**
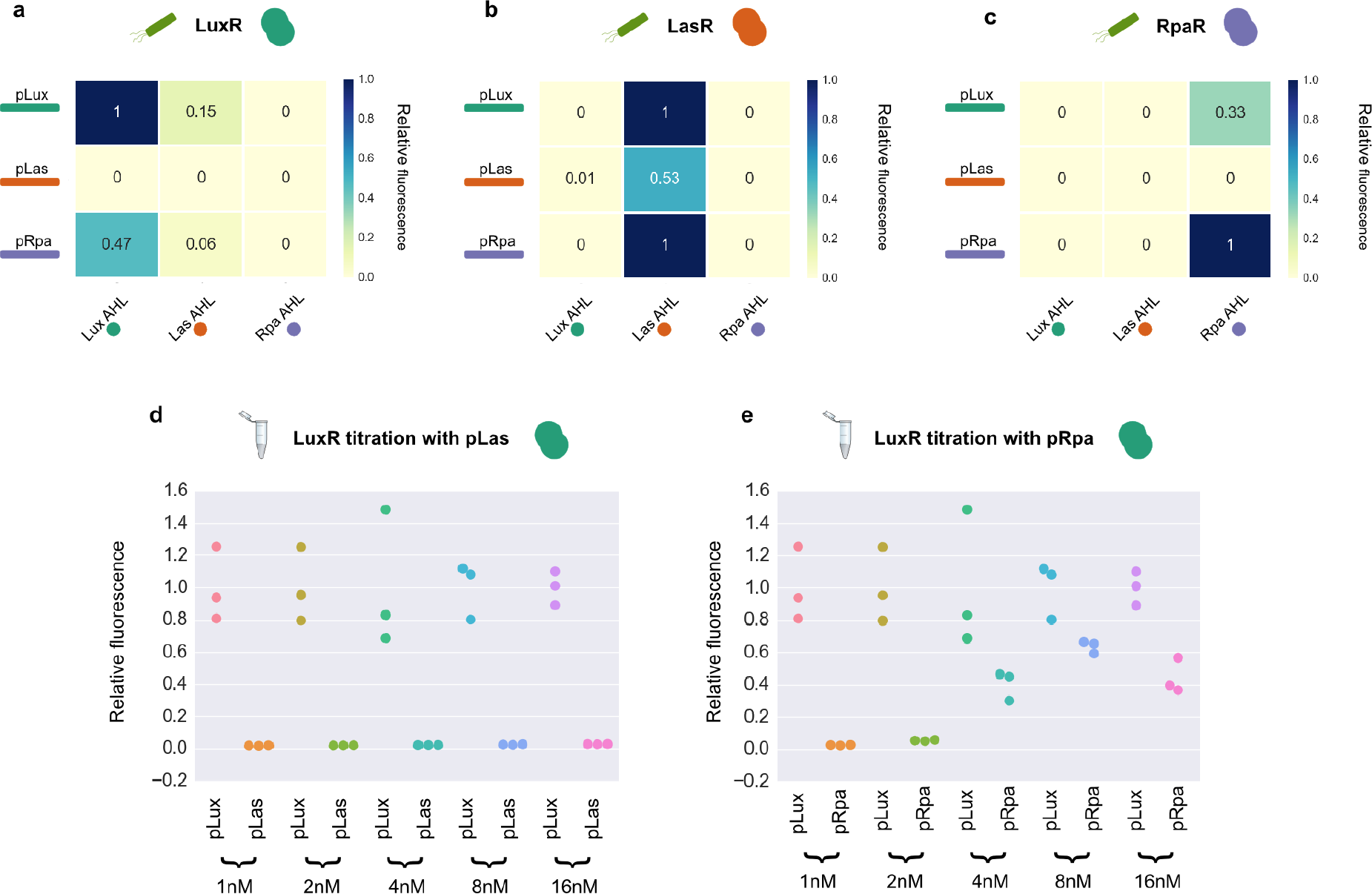
*in vivo* crosstalk and TX-TL titration experiments. (A) LuxR crosstalk heatmap *in vivo* at 1uM of AHL. Relative fluorescence is calculated for each combination as the average of the 95^th^percentile fluorescence, normalized by OD600, value across four replicates divided by the maximum observed across all nine combinations. (B) LasR crosstalk heatmap. (C) RpaR crosstalk heatmap. (D) LuxR titration with pLas at 1uM AHL. Relative fluorescence calculated at each concentration of LuxR DNA (1-16nM) as 95^th^ percentile fluorescence / mean of three pLux-GFP 95^th^ percentile fluorescence at the same concentration of LuxR. (E) LuxR titration with pRpa.

A possible explanation for the increased genetic crosstalk *in vivo* is a higher ratio of transcription factor to promoter DNA, increasing the amount of off-target activation. If this change in ratio caused our observed crosstalk pattern *in vivo*, we should be able to reproduce the pattern in TX-TL by increasing the amount of DNA encoding for the transcription factor while holding the amount of promoter DNA constant. We chose to test this hypothesis with the LuxR system. We performed titration experiments holding the concentration of each promoter constant, while increasing the LuxR concentration from 1nM DNA up to 16nM of DNA. As the amount of LuxR DNA was increased the relative expression from pRpa increased while the pLas promoter remained quiescent (Figure 2D-E). This matches the *in vivo* result that LuxR activates both pLux and pRpa, but not pLas. This suggests that one factor that could explain the discrepancy between cell-free and *in vivo* characterization is differing effective concentrations of transcription factor relative to free promoter.

Cell-free experiments accurately predicted both signal level crosstalk mediated by improper AHL-TF binding, and genetic crosstalk caused by off-target TF-DNA binding and recruitment of RNA polymerase. The crosstalk patterns elucidated are not only useful for designing population-level genetic circuits, but also provide a template for using cell-free systems to characterize quorum sensing parts.

As microbiologists continue to identify new quorum sensing systems, rapid prototyping to map their crosstalk interactions will become increasingly valuable to synthetic biologists. Using cell-free platforms to rapidly characterize quorum system interaction spaces will allow for high-throughput testing across large design spaces. Exciting future work in charactering quorum sensing systems will include characterization of the remaining circuit component, AHL synthases, using both cell-free and *in vivo* systems.

## Materials and methods

### TX-TL

TX-TL was prepared as previously described.^9^ All experiments were conducted at 5uL total volume. Plates were read in a BioTek Synergy H1M with excitation wavelength of 485nm and emission wavelength of 515nm for deGFP.

### In vivo

JM109 cells were transformed with both a plasmid constitutively expressing activator plasmid (p15a chloramphenicol) and a plasmid with the AHL promoter regulating GFP expression (colE1 kanamycin). Three colonies from each plate were inoculated into LB containing chloramphenicol (34ug/mL) and kanamycin (50ug/mL) overnight. Cultures were then diluted 1:100 fold and grown until mid-log phase (0.3-0.6 OD600). Cultures were then diluted into M9 media containing 1% glucose and 1uM of the appropriate AHL, or 0.1% DMSO as a negative control. Cells were grown in 96 well plates in an BioTek Synergy H1M. Cells were grown at 37°C.

## Data Analysis

Data analysis was performed using custom Python scripts. All raw data and code used in this manuscript is available at github.com/adhalleran/QS. All sequence files for constructs used in this manuscript is also available at github.com/adhalleran/QS.

## Acknowledgements

The authors would like to thank Sam Clamons, Andrey Shur and Vipul Singh for fruitful discussions. We also thank John Marken for helpful comments on the manuscript. Plasmid vectors were provided by Douglas Densmore at the Cross-disciplinary Integration of Design Automation Research lab (Addgene Kit # 1000000059). The project depicted was sponsored by the Defense Advanced Research Projects Agency (Agreement HR0011-17-2-0008). The content of the information does not necessarily reflect the position or the policy of the Government, and no official endorsement should be inferred.

## Supporting Information

**Supporting Figure 1:**
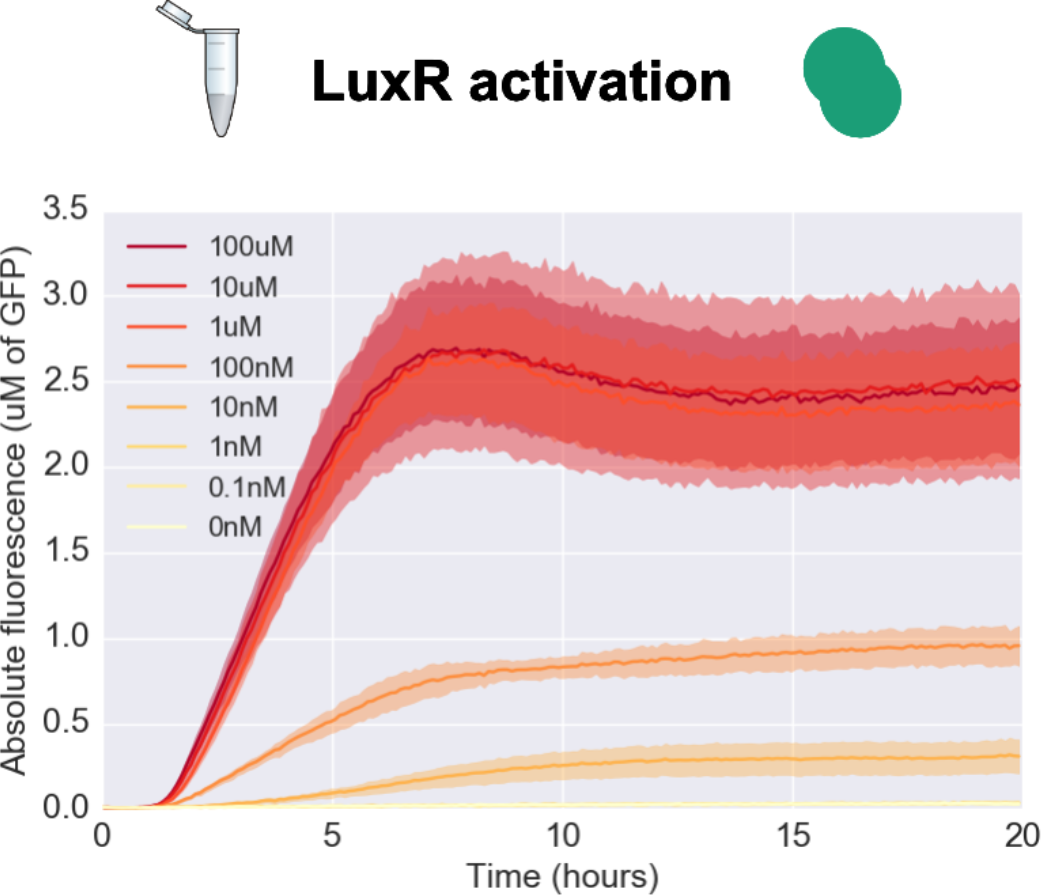
TX-TL time traces from LuxR (2nM DNA) + pLux-GFP (4nM DNA) across a range of Lux AHL concentrations. Solid lines represent the mean of four replicates, shading represents +/- one standard deviation.

**Supporting Figure 2:**
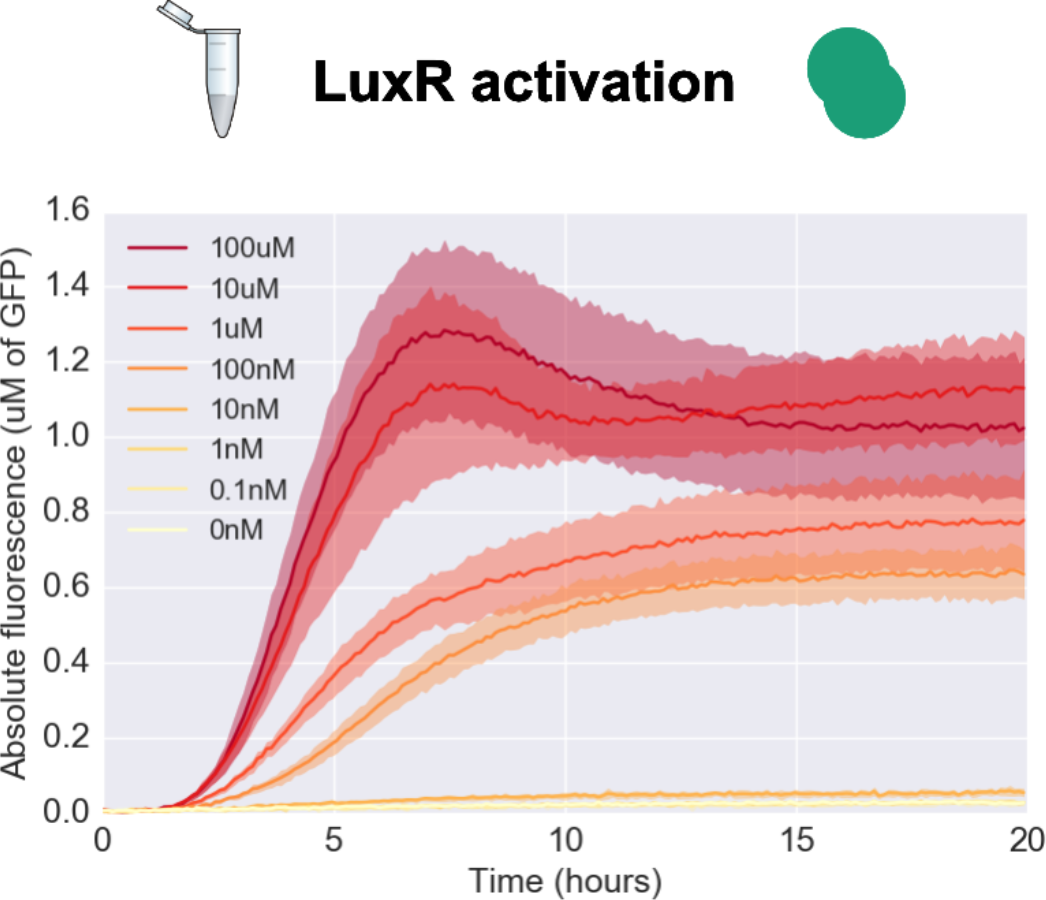
TX-TL time traces from LuxR (2nM DNA) + pLux-GFP (4nM DNA) across a range of Las AHL concentrations. Solid lines represent the mean of four replicates, shading represents +/- one standard deviation.

**Supporting Figure 3:**
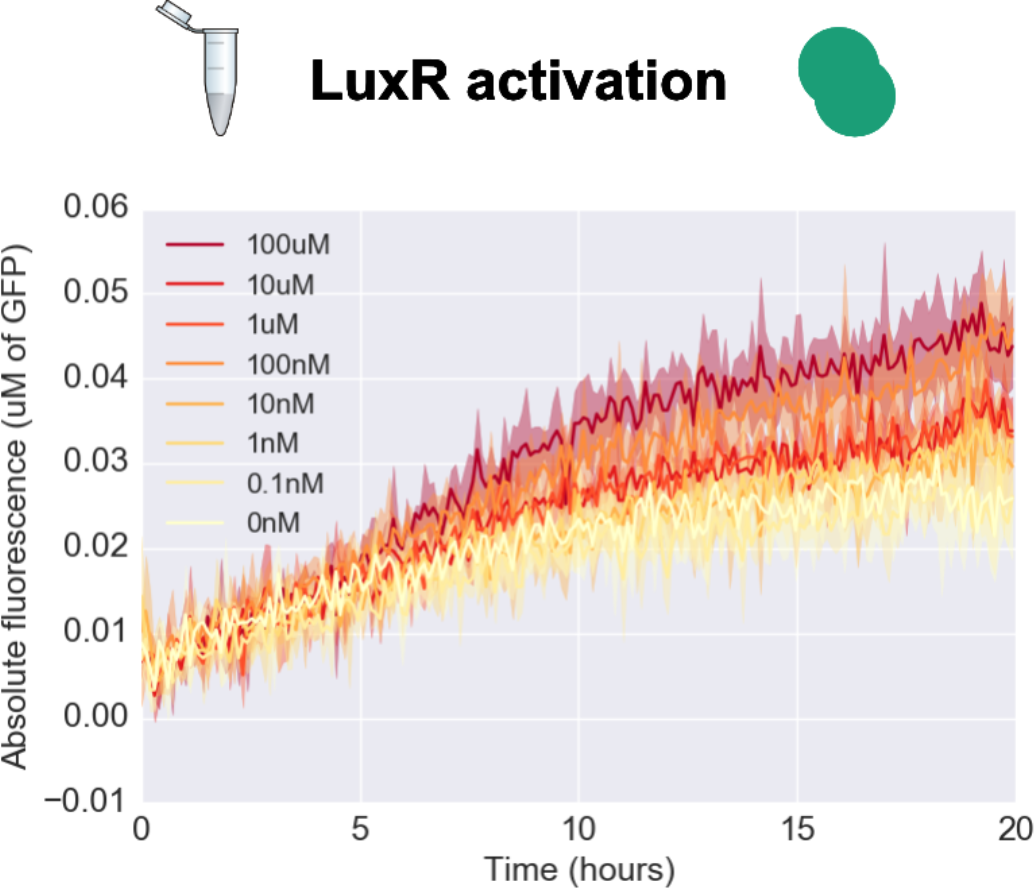
TX-TL time traces from LuxR (2nM DNA) + pLux-GFP (4nM DNA) across a range of Rpa AHL concentrations. Solid lines represent the mean of four replicates, shading represents +/- one standard deviation.

**Supporting Figure 4:**
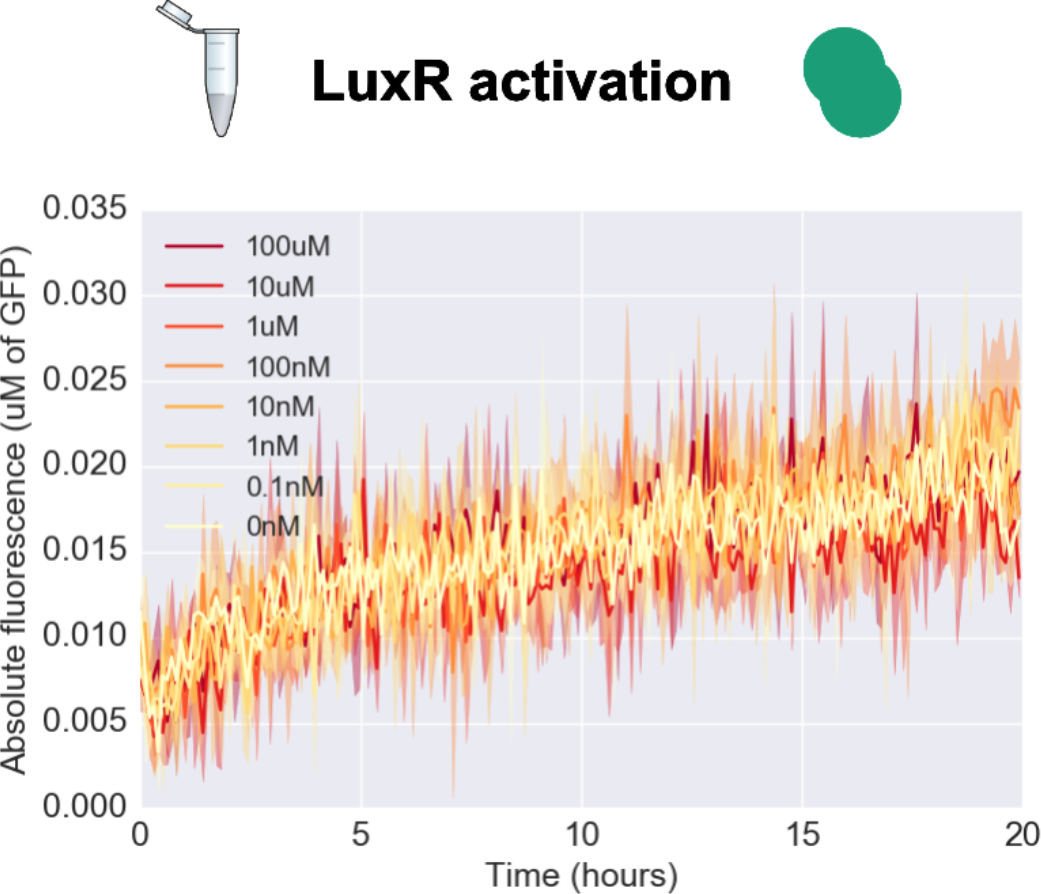
TX-TL time traces from LuxR (2nM DNA) + pLas-GFP (4nM DNA) across a range of Lux AHL concentrations. Solid lines represent the mean of four replicates, shading represents +/- one standard deviation.

**Supporting Figure 5:**
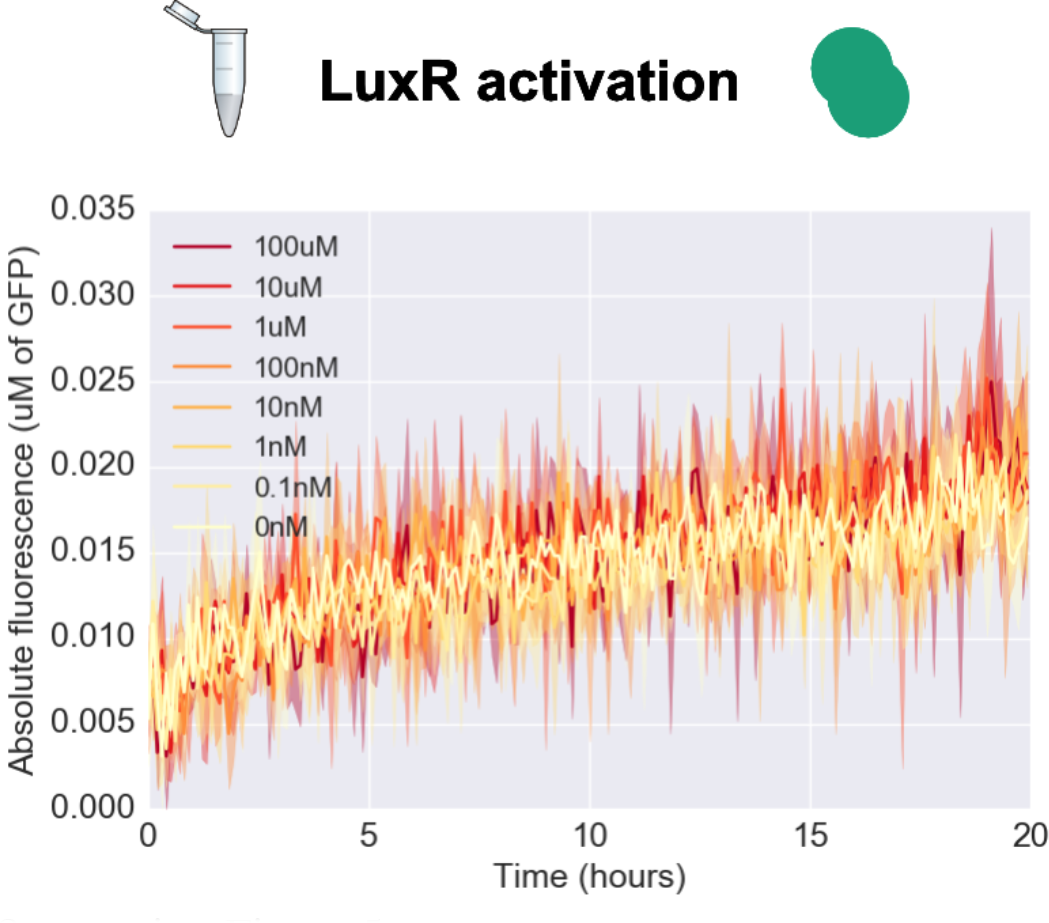
TX-TL time traces from LuxR (2nM DNA) + pLas-GFP (4nM DNA) across a range of Las AHL concentrations. Solid lines represent the mean of four replicates, shading represents +/- one standard deviation.

**Supporting Figure 6:**
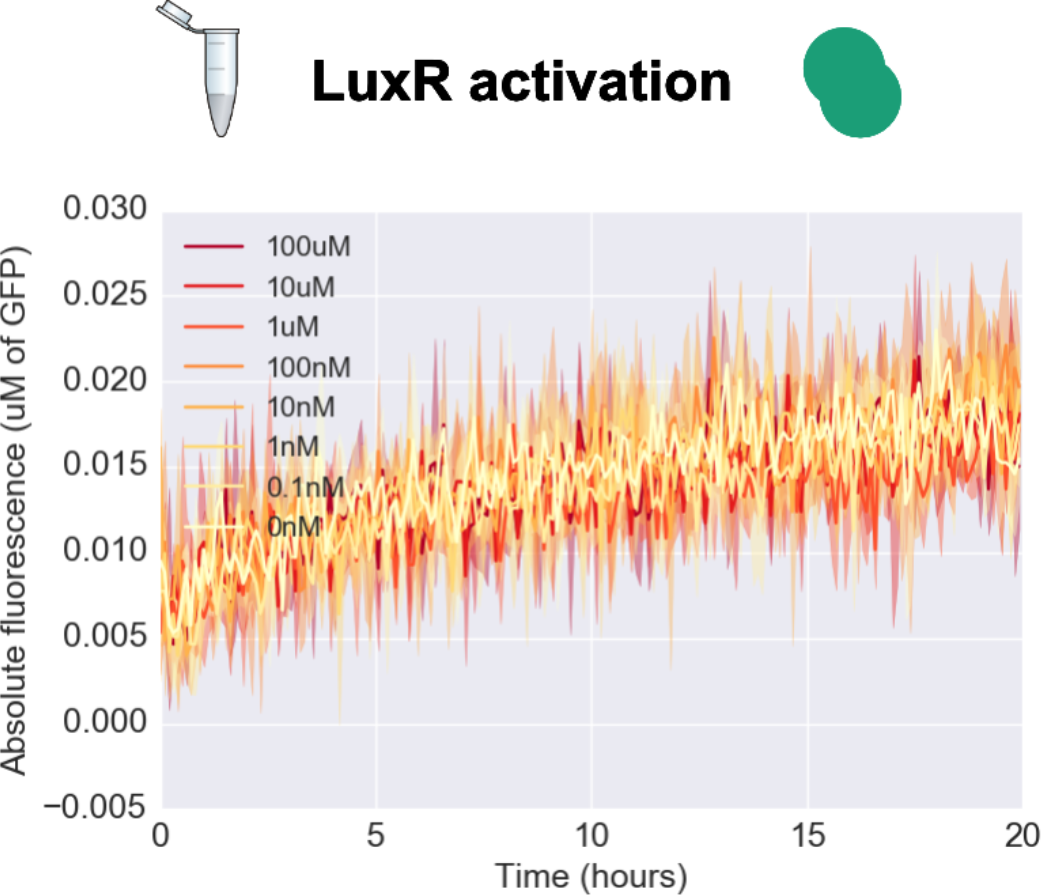
TX-TL time traces from LuxR (2nM DNA) + pLas-GFP (4nM DNA) across a range of Rpa AHL concentrations. Solid lines represent the mean of four replicates, shading represents +/- one standard deviation.

**Supporting Figure 7:**
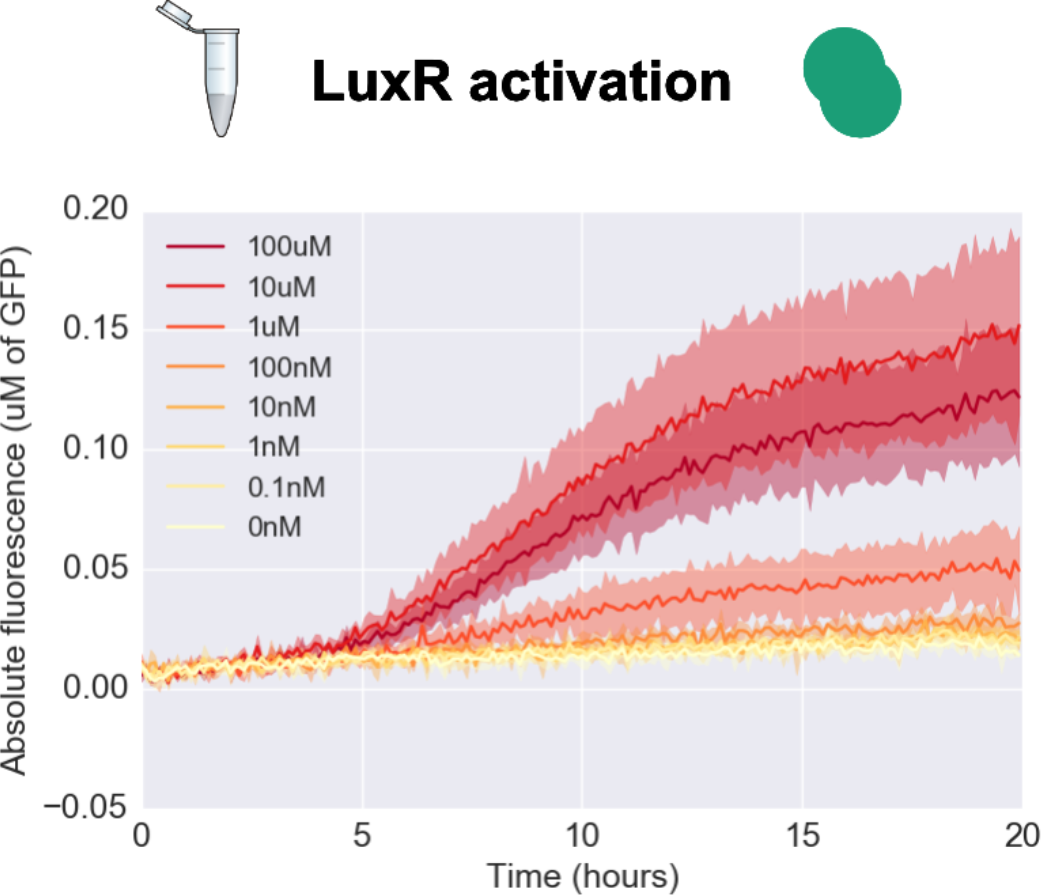
TX-TL time traces from LuxR (2nM DNA) + pRpa-GFP (4nM DNA) across a range of Lux AHL concentrations. Solid lines represent the mean of four replicates, shading represents +/- one standard deviation.

**Supporting Figure 8:**
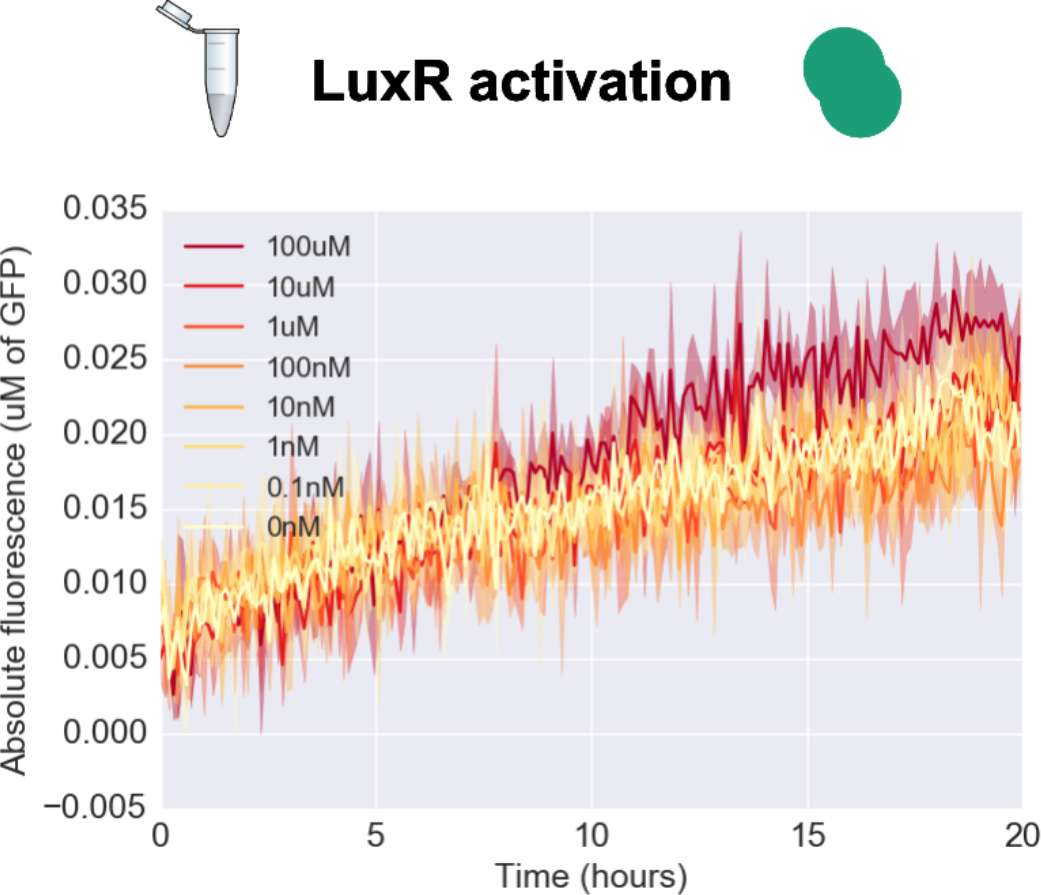
TX-TL time traces from LuxR (2nM DNA) + pRpa-GFP (4nM DNA) across a range of Las AHL concentrations. Solid lines represent the mean of four replicates, shading represents +/- one standard deviation.

**Supporting Figure 9:**
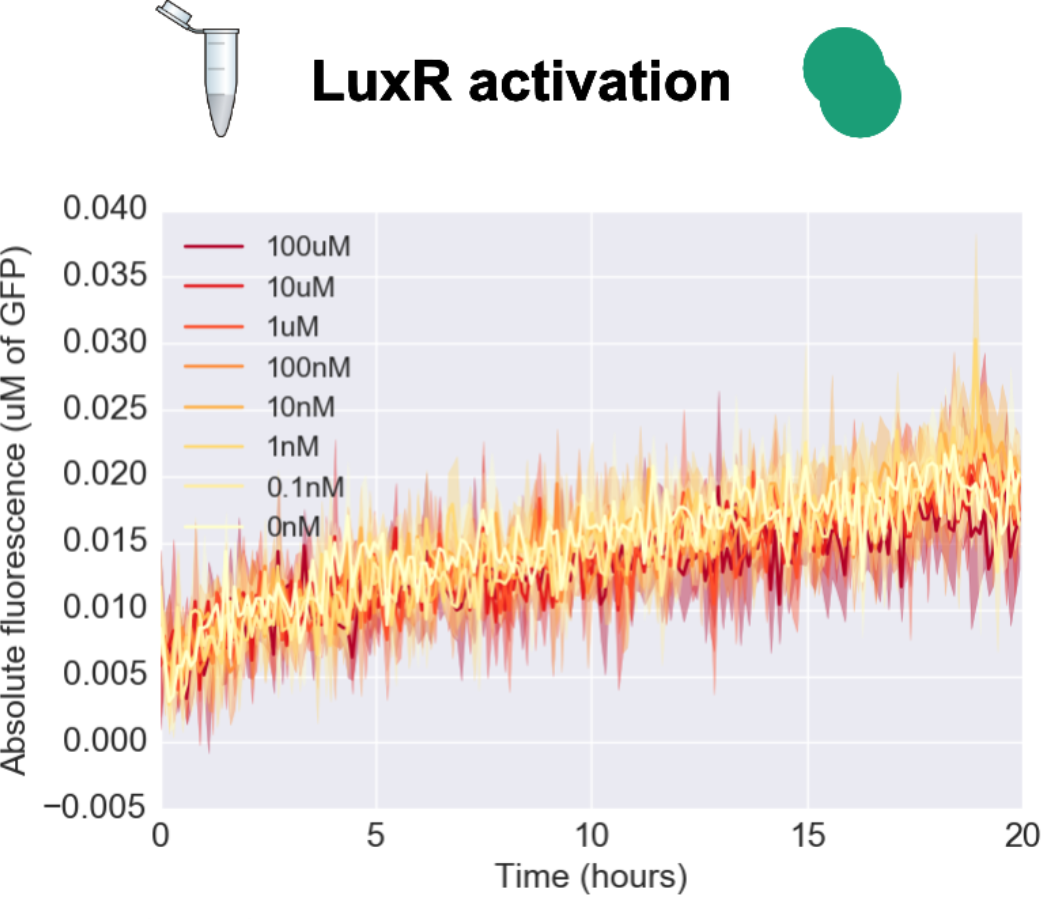
TX-TL time traces from LuxR (2nM DNA) + pRpa-GFP (4nM DNA) across a range of Rpa AHL concentrations. Solid lines represent the mean of four replicates, shading represents +/- one standard deviation.

**Supporting Figure 10:**
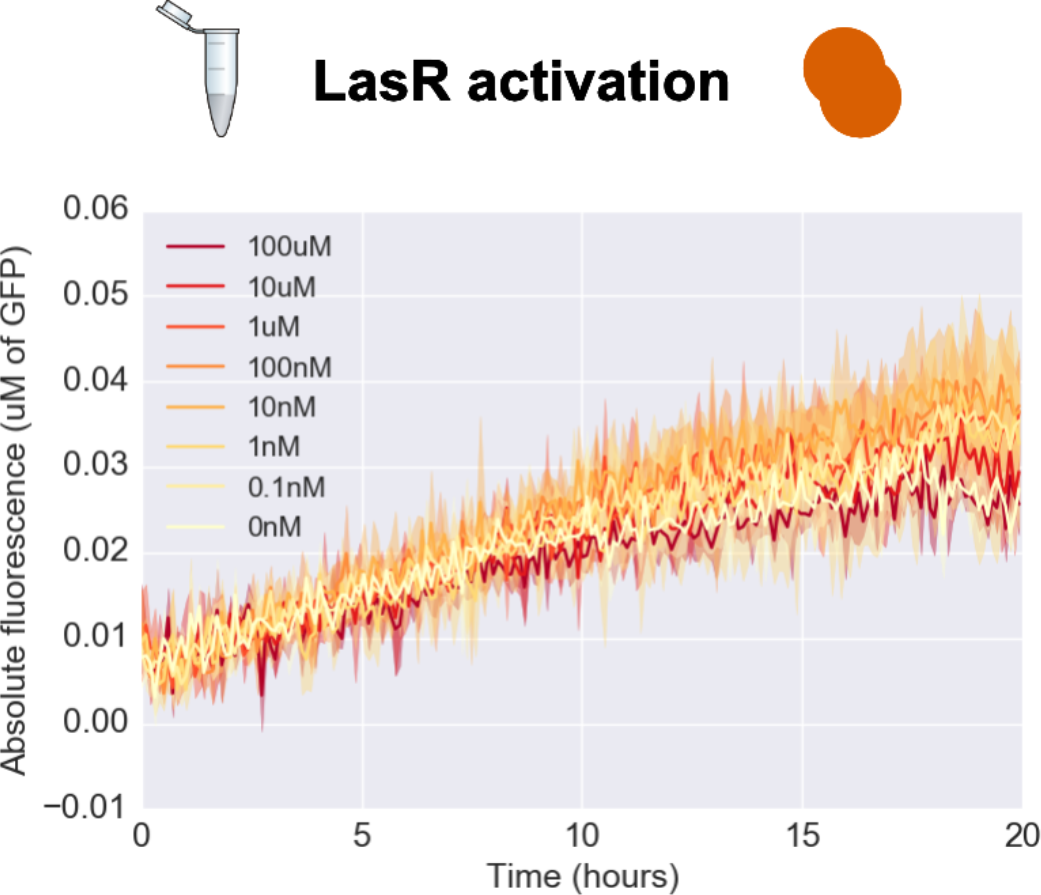
TX-TL time traces from LasR (2nM DNA) + pLux-GFP (4nM DNA) across a range of Lux AHL concentrations. Solid lines represent the mean of four replicates, shading represents +/- one standard deviation.

**Supporting Figure 11:**
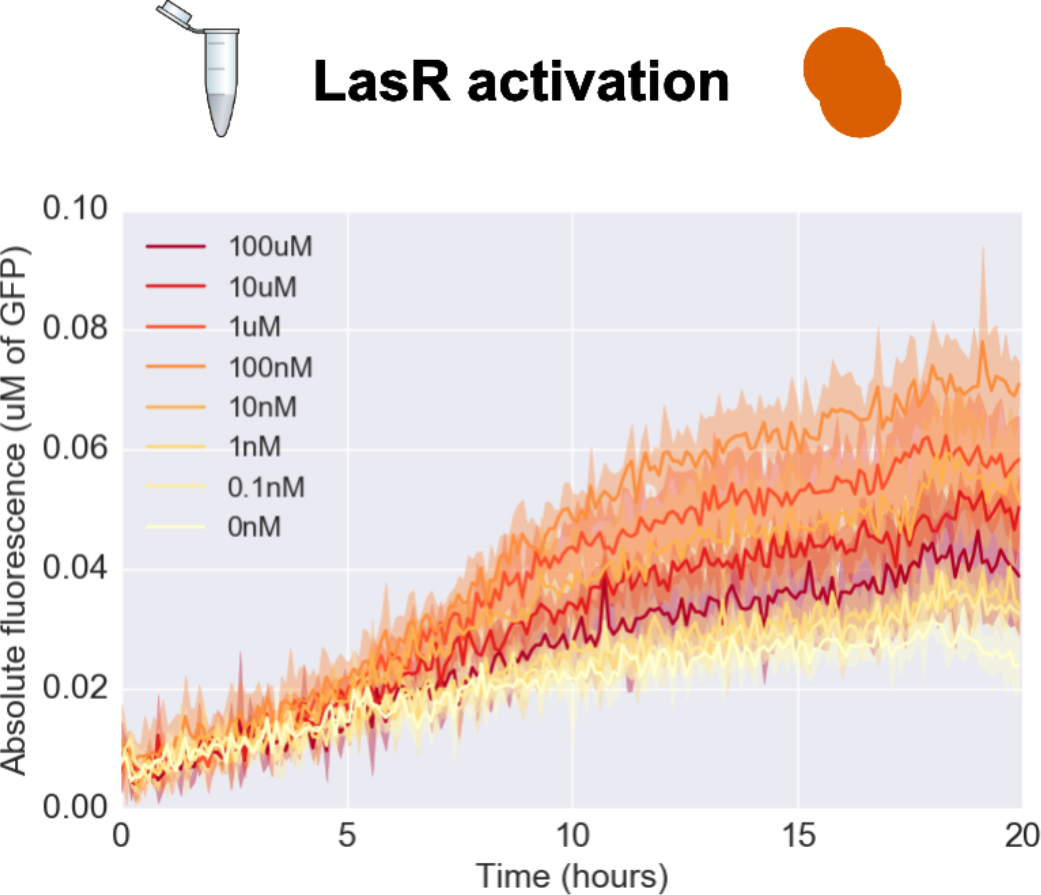
TX-TL time traces from LasR (2nM DNA) + pLux-GFP (4nM DNA) across a range of Las AHL concentrations. Solid lines represent the mean of four replicates, shading represents +/- one standard deviation.

**Supporting Figure 12:**
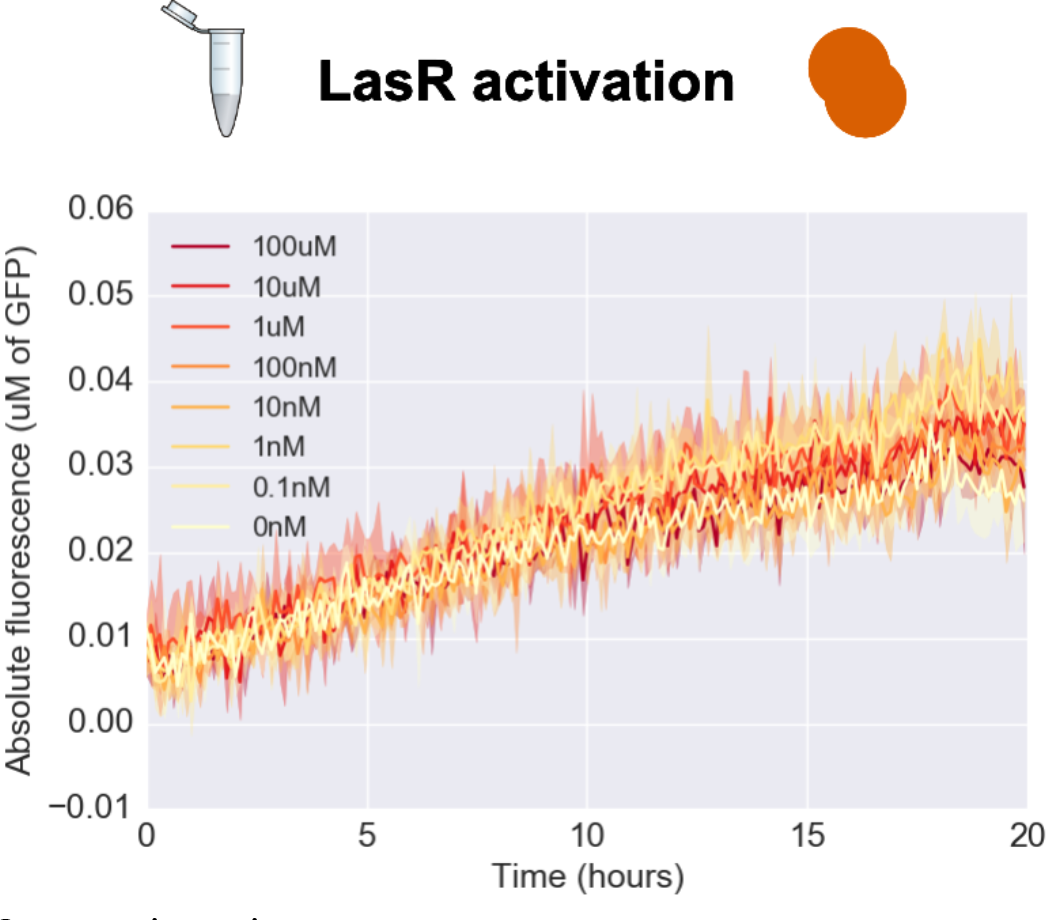
TX-TL time traces from LasR (2nM DNA) + pLux-GFP (4nM DNA) across a range of Rpa AHL concentrations. Solid lines represent the mean of four replicates, shading represents +/- one standard deviation.

**Supporting Figure 13:**
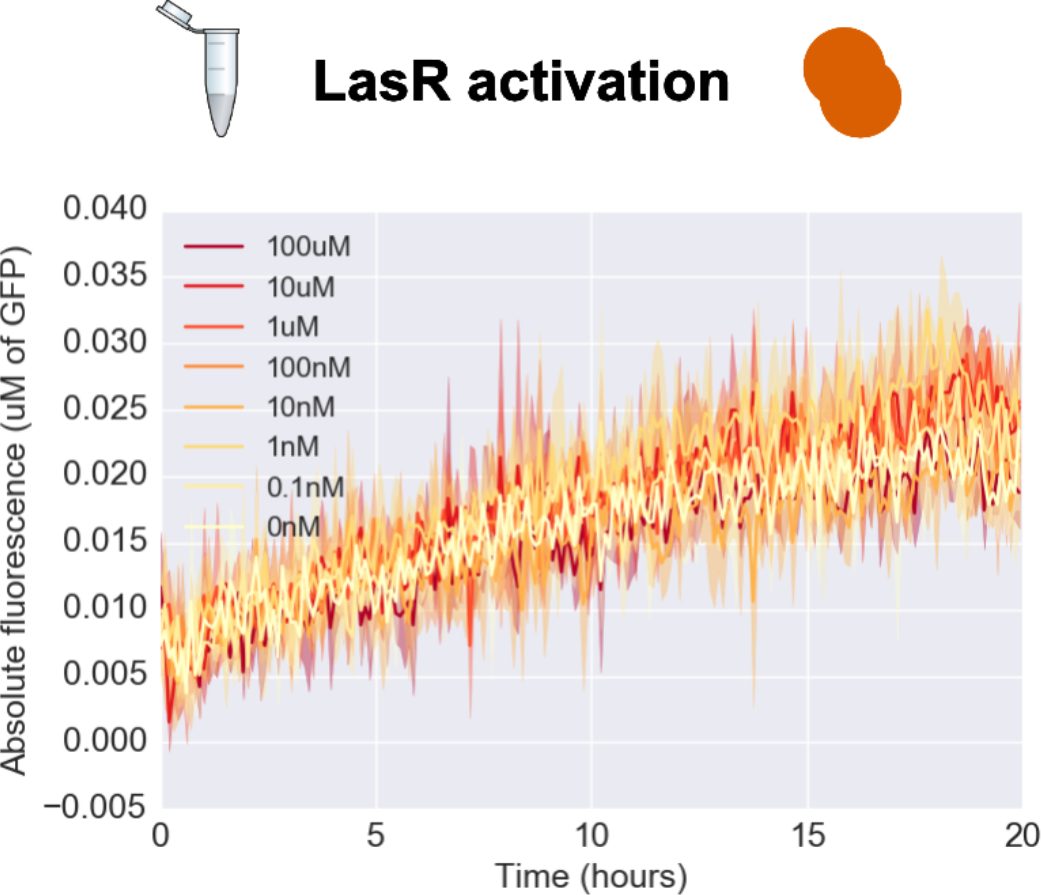
TX-TL time traces from LasR (2nM DNA) + pLas-GFP (4nM DNA) across a range of Lux AHL concentrations. Solid lines represent the mean of four replicates, shading represents +/- one standard deviation.

**Supporting Figure 14:**
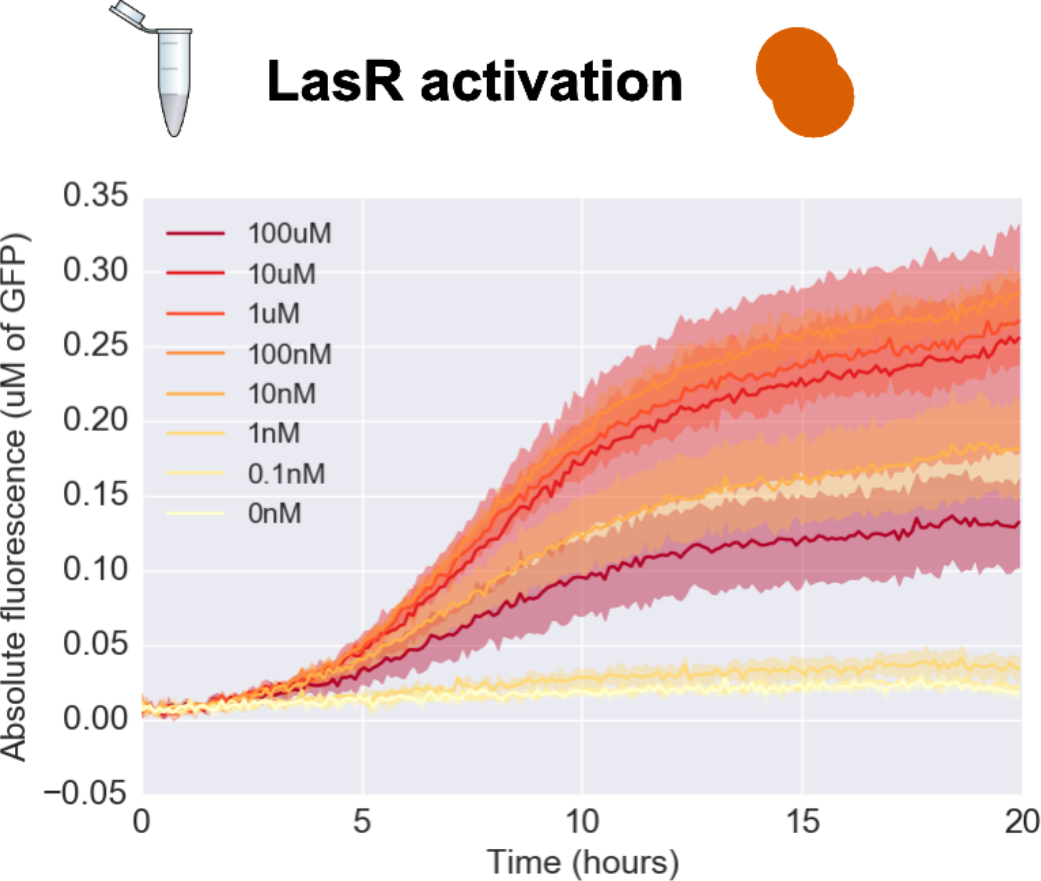
TX-TL time traces from LasR (2nM DNA) + pLas-GFP (4nM DNA) across a range of Las AHL concentrations. Solid lines represent the mean of four replicates, shading represents +/- one standard deviation.

**Supporting Figure 15:**
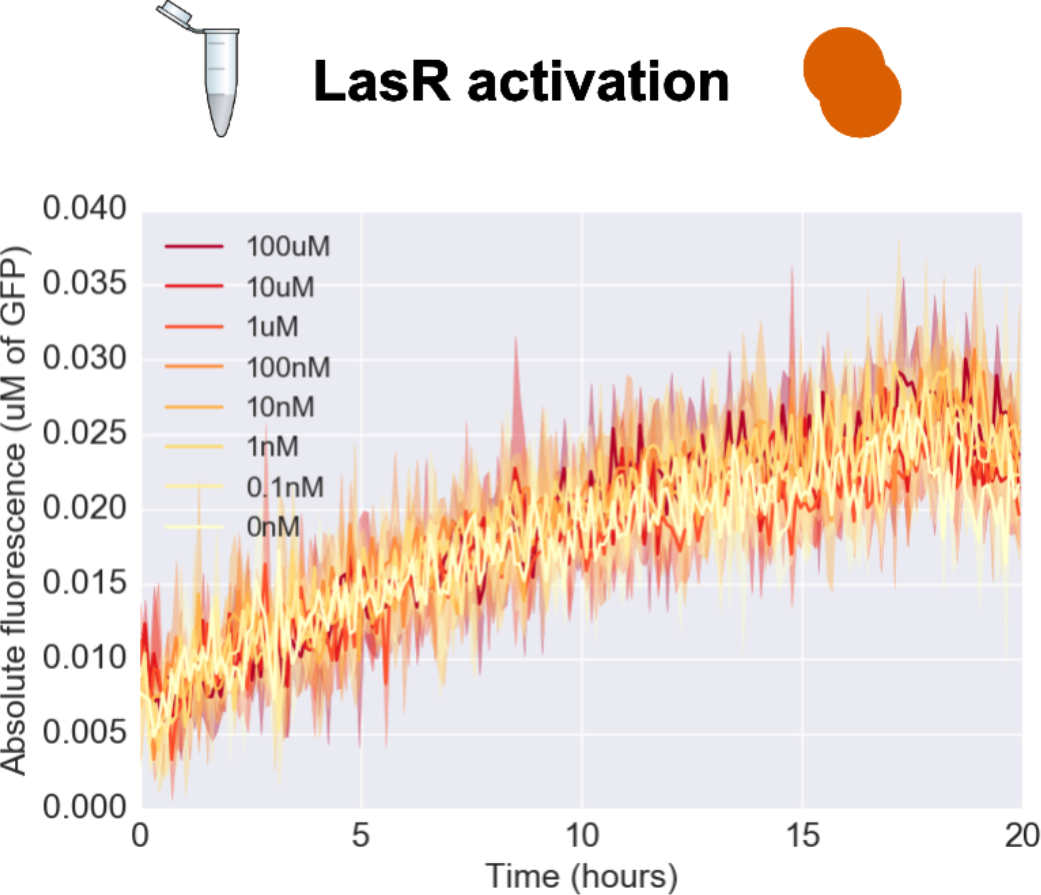
TX-TL time traces from LasR (2nM DNA) + pLas-GFP (4nM DNA) across a range of Rpa AHL concentrations. Solid lines represent the mean of four replicates, shading represents +/- one standard deviation.

**Supporting Figure 16:**
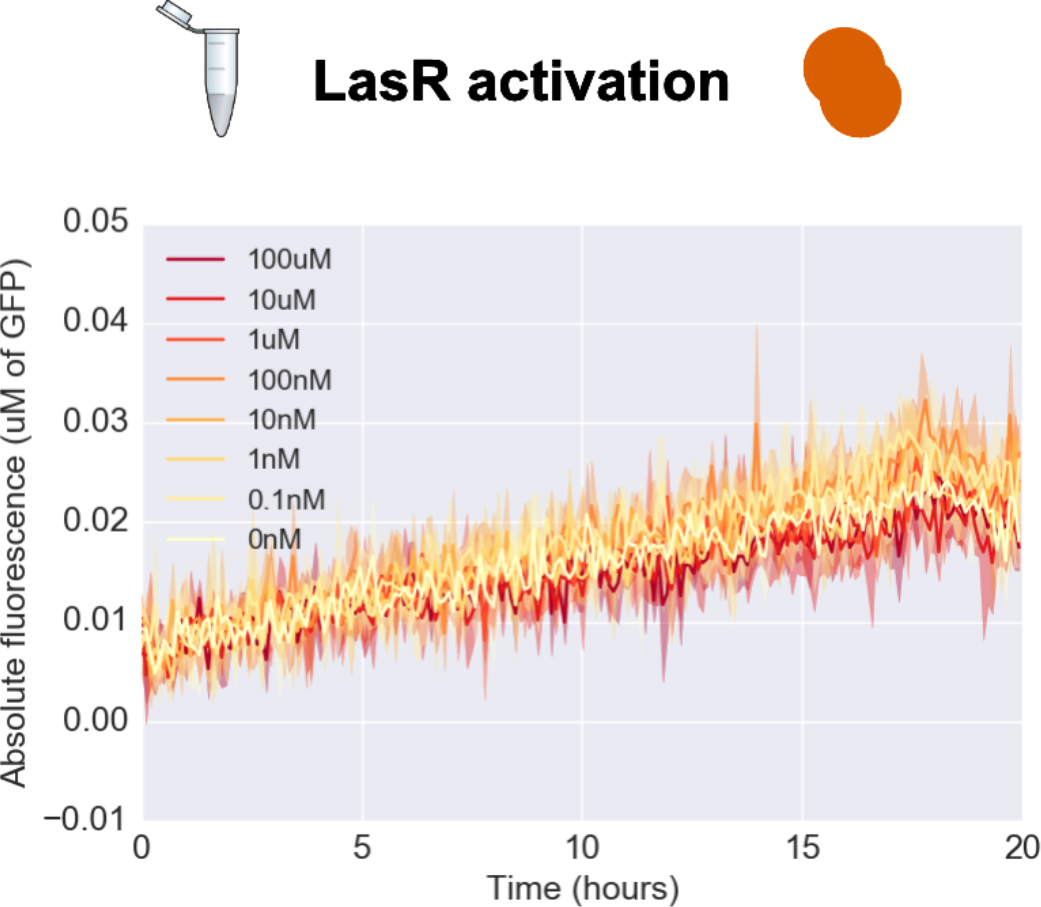
TX-TL time traces from LasR (2nM DNA) + pRpa-GFP (4nM DNA) across a range of Lux AHL concentrations. Solid lines represent the mean of four replicates, shading represents +/- one standard deviation.

**Supporting Figure 17:**
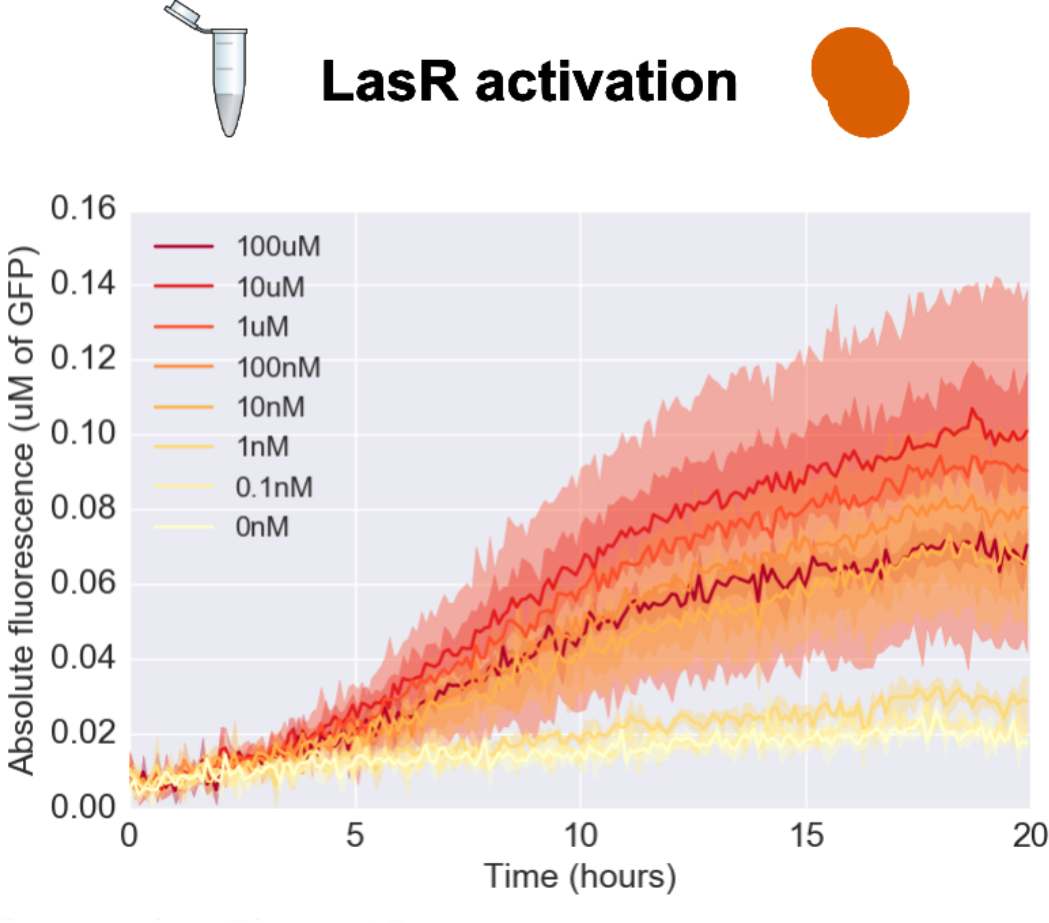
TX-TL time traces from LasR (2nM DNA) + pRpa-GFP (4nM DNA) across a range of Las AHL concentrations. Solid lines represent the mean of four replicates, shading represents +/- one standard deviation.

**Supporting Figure 18:**
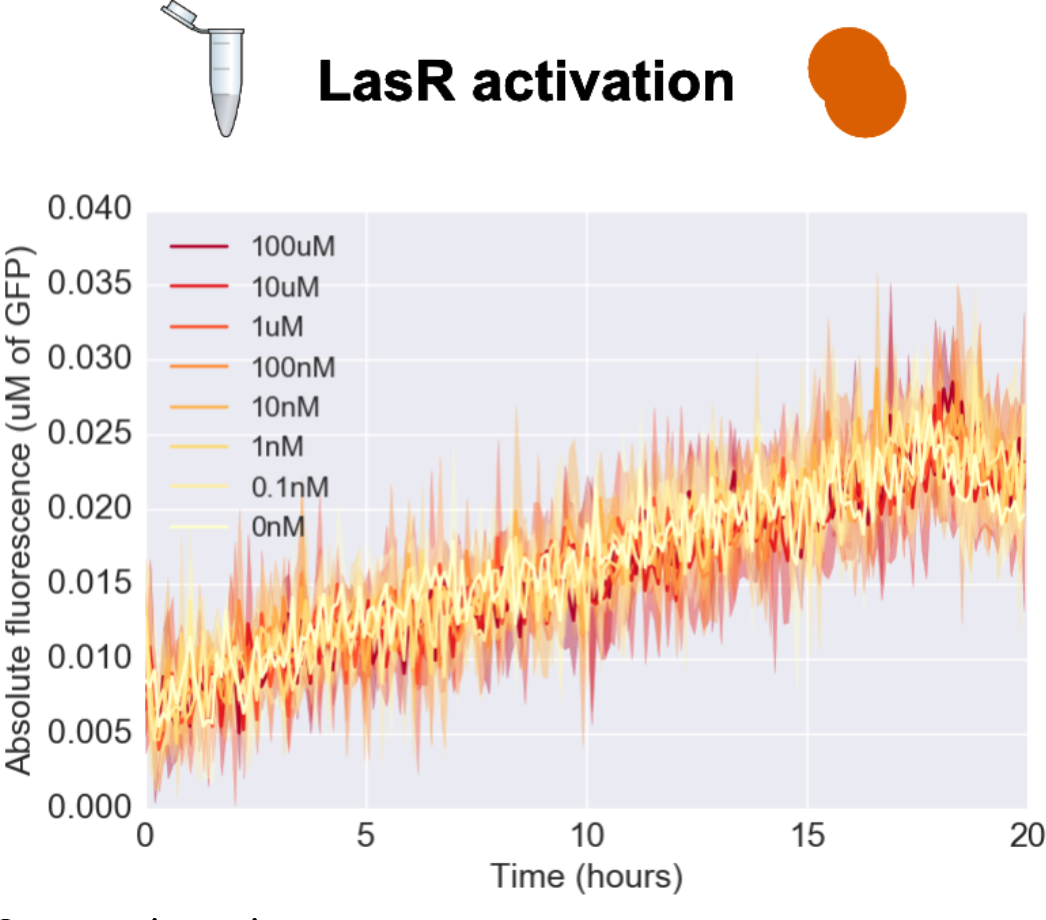
TX-TL time traces from LasR (2nM DNA) + pRpa-GFP (4nM DNA) across a range of Rpa AHL concentrations. Solid lines represent the mean of four replicates, shading represents +/- one standard deviation.

**Supporting Figure 19:**
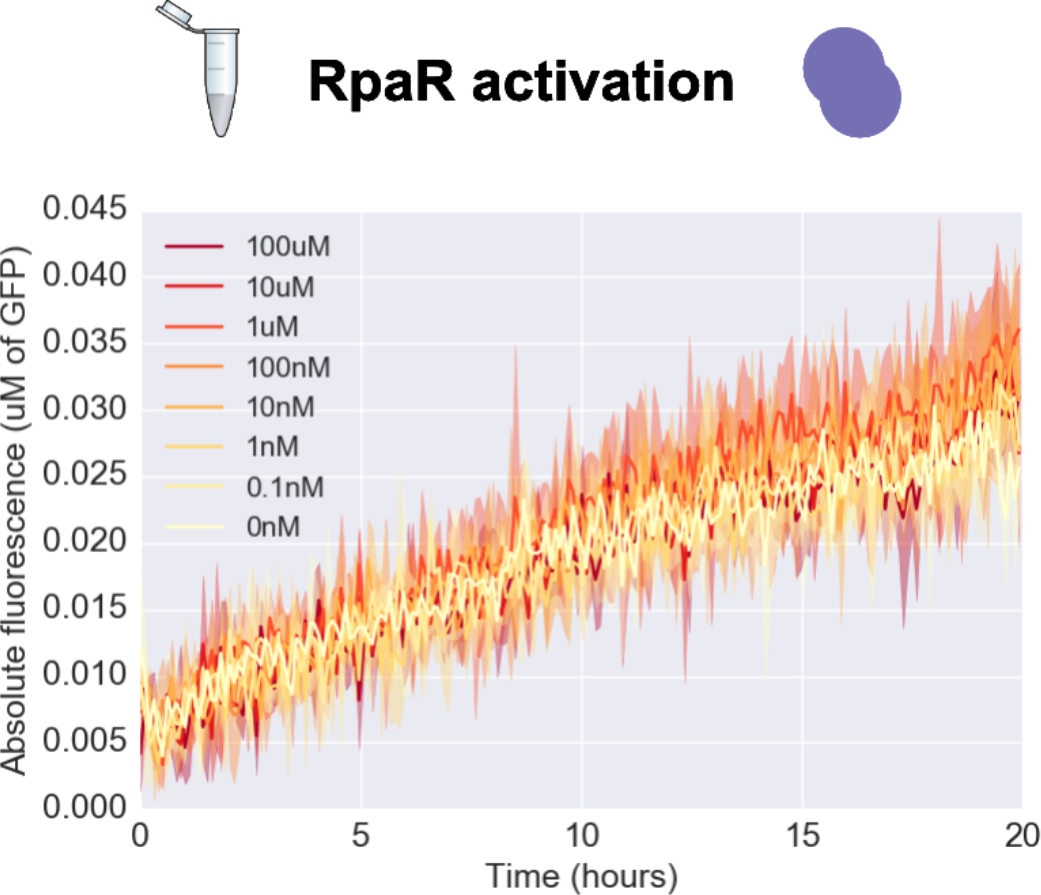
TX-TL time traces from RpaR (2nM DNA) + pLux-GFP (4nM DNA) across a range of Lux AHL concentrations. Solid lines represent the mean of four replicates, shading represents +/- one standard deviation.

**Supporting Figure 20:**
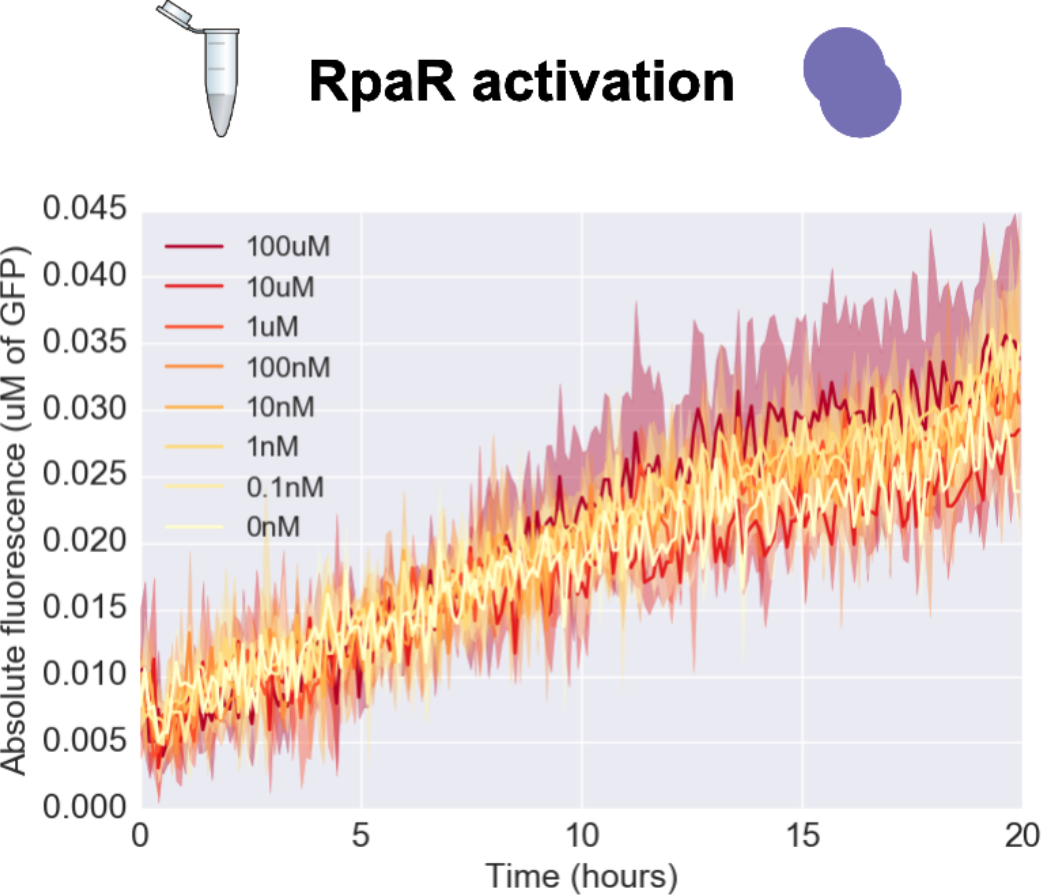
TX-TL time traces from RpaR (2nM DNA) + pLux-GFP (4nM DNA) across a range of Las AHL concentrations. Solid lines represent the mean of four replicates, shading represents +/- one standard deviation.

**Supporting Figure 21:**
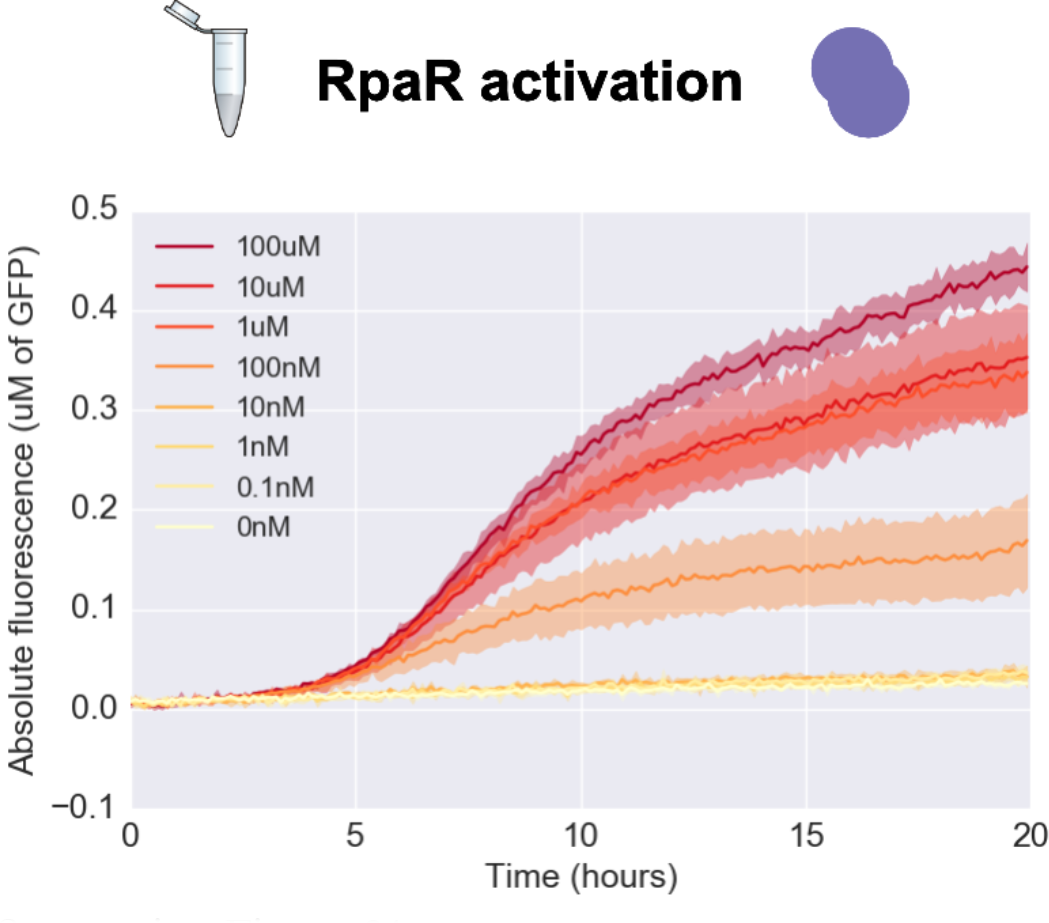
TX-TL time traces from RpaR (2nM DNA) + pLux-GFP (4nM DNA) across a range of Rpa AHL concentrations. Solid lines represent the mean of four replicates, shading represents +/- one standard deviation.

**Supporting Figure 22:**
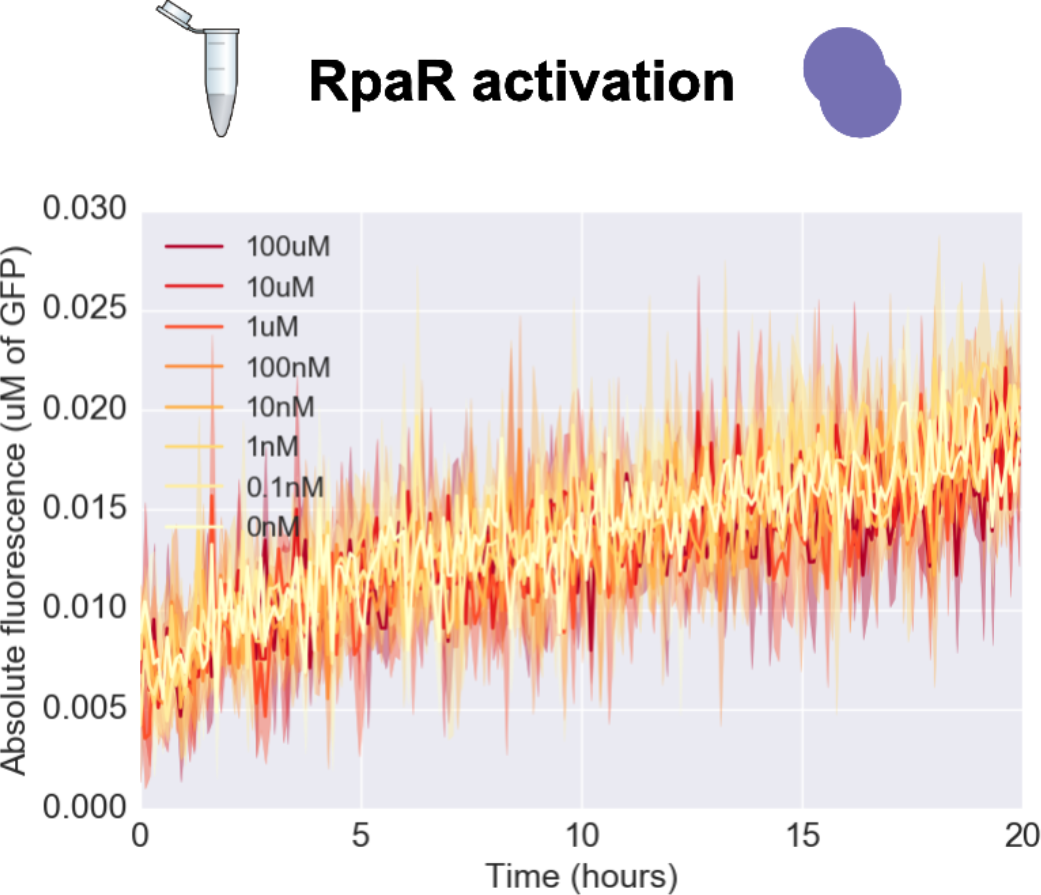
TX-TL time traces from RpaR (2nM DNA) + pLas-GFP (4nM DNA) across a range of Lux AHL concentrations. Solid lines represent the mean of four replicates, shading represents +/- one standard deviation.

**Supporting Figure 23:**
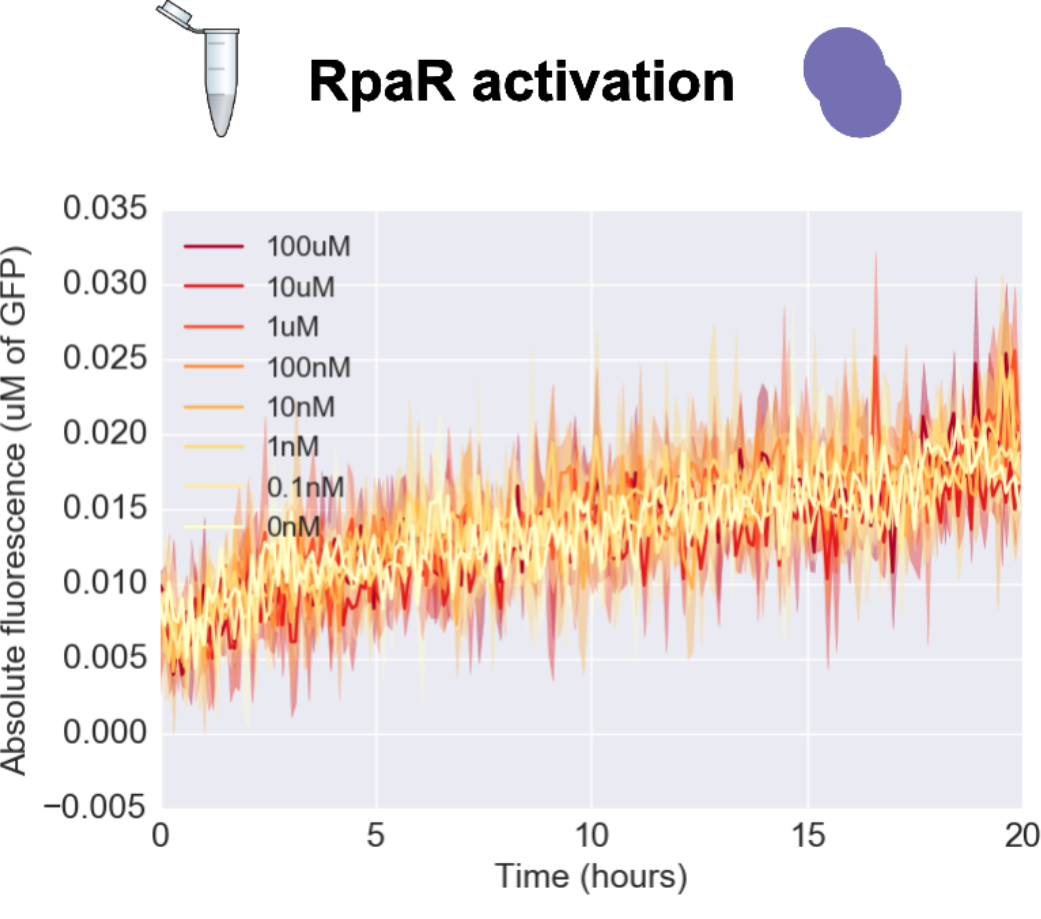
TX-TL time traces from RpaR (2nM DNA) + pLas-GFP (4nM DNA) across a range of Las AHL concentrations. Solid lines represent the mean of four replicates, shading represents +/- one standard deviation.

**Supporting Figure 24:**
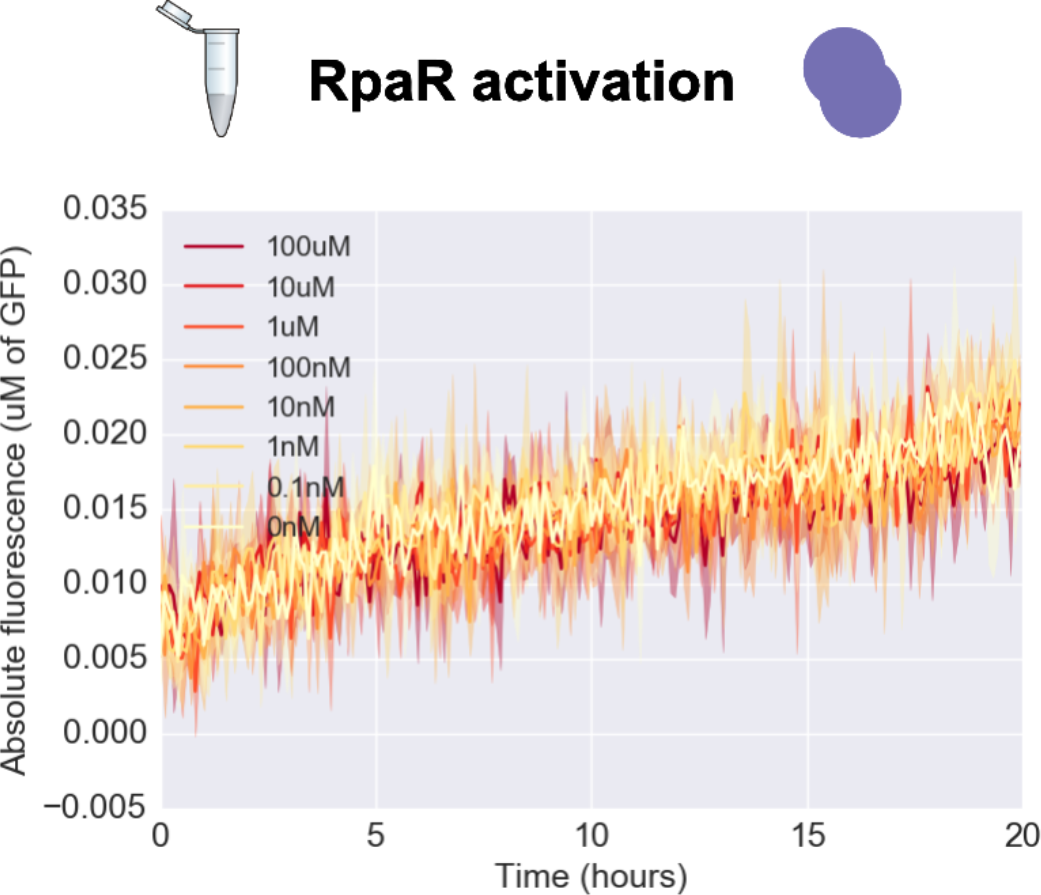
TX-TL time traces from RpaR (2nM DNA) + pLas-GFP (4nM DNA) across a range of Rpa AHL concentrations. Solid lines represent the mean of four replicates, shading represents +/- one standard deviation.

**Supporting Figure 25:**
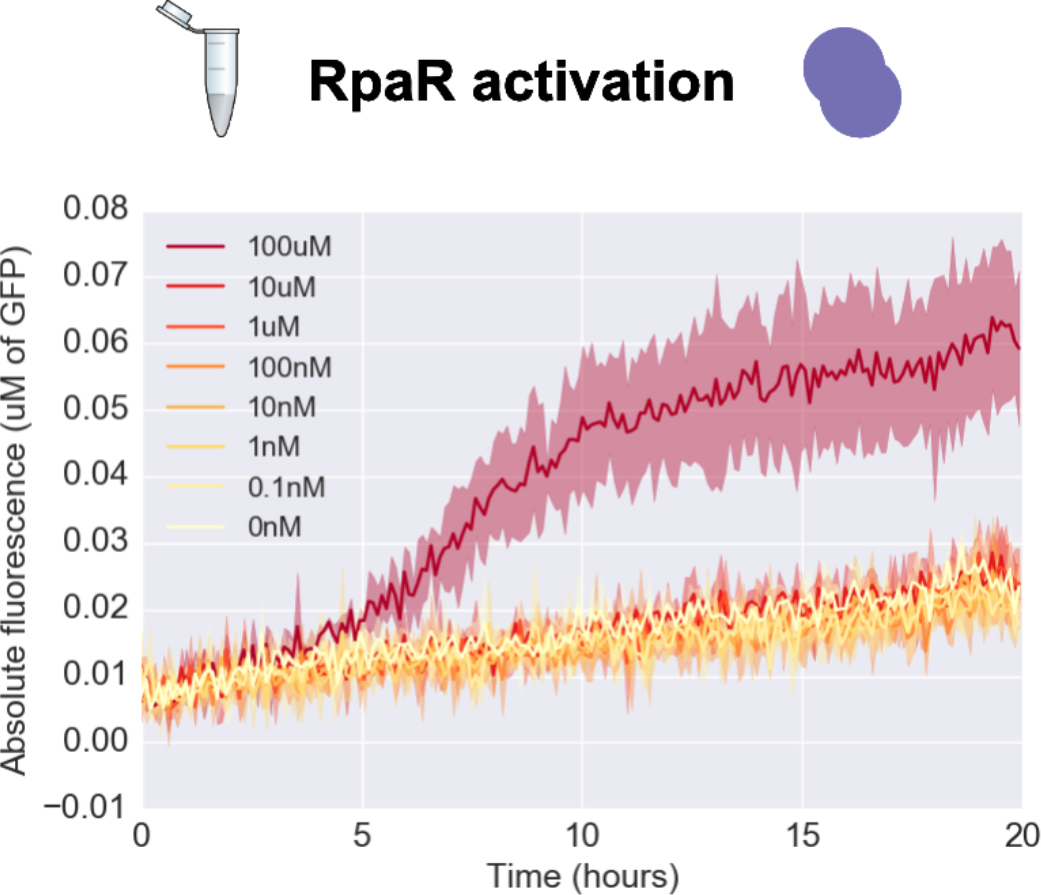
TX-TL time traces from RpaR (2nM DNA) + pRpa-GFP (4nM DNA) across a range of Lux AHL concentrations. Solid lines represent the mean of four replicates, shading represents +/- one standard deviation.

**Supporting Figure 26:**
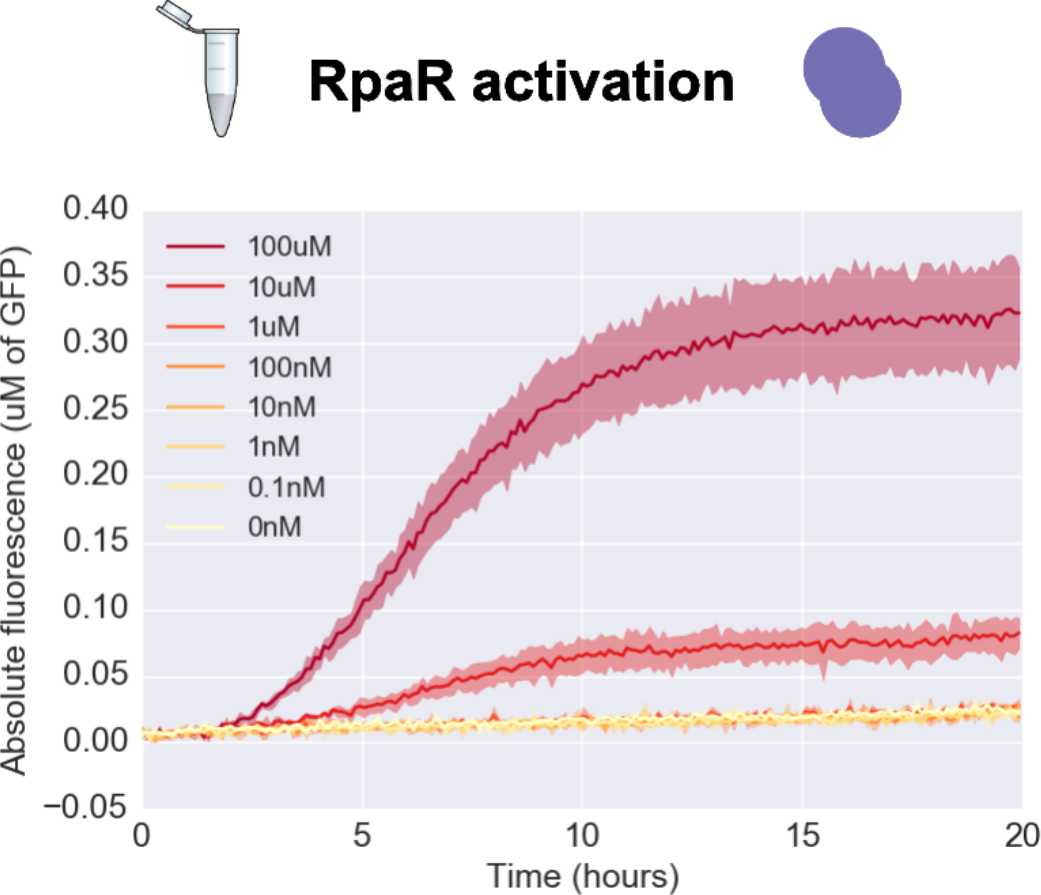
TX-TL time traces from RpaR (2nM DNA) + pRpa-GFP (4nM DNA) across a range of Las AHL concentrations. Solid lines represent the mean of four replicates, shading represents +/- one standard deviation.

**Supporting Figure 27:**
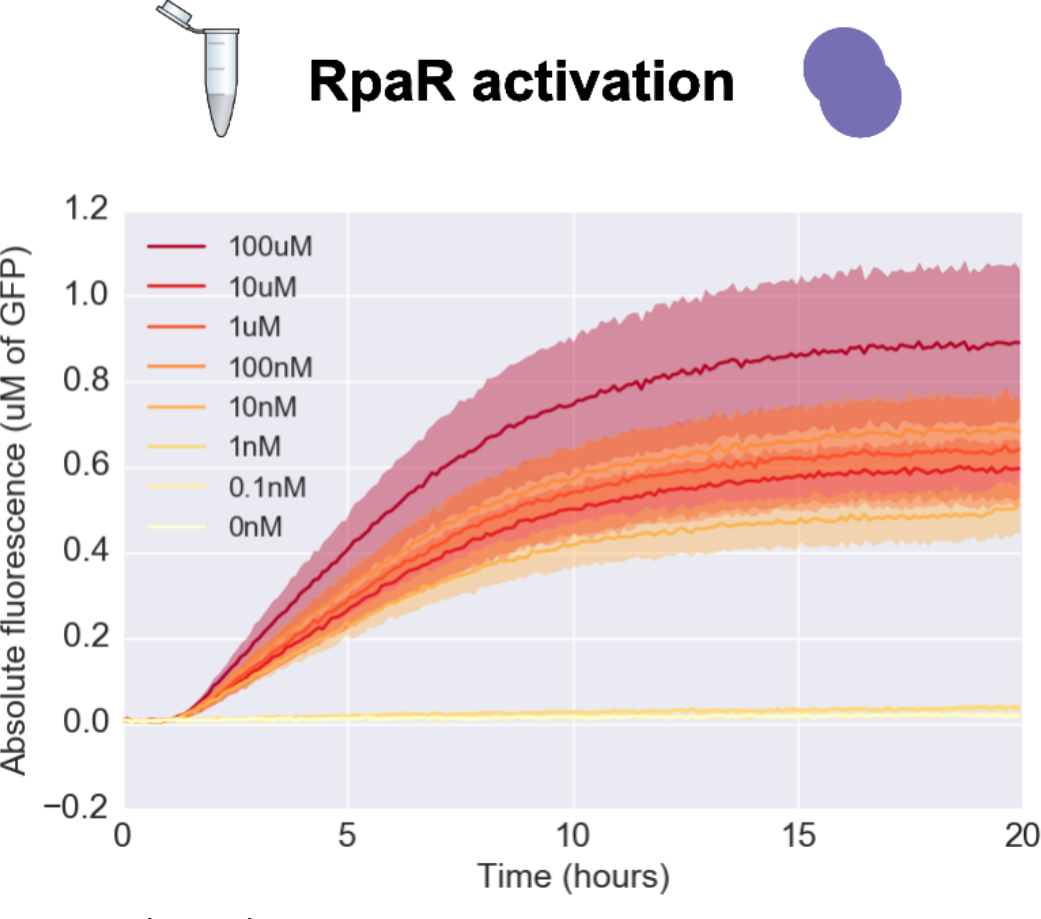
TX-TL time traces from RpaR (2nM DNA) + pRpa-GFP (4nM DNA) across a range of Rpa AHL concentrations. Solid lines represent the mean of four replicates, shading represents +/- one standard deviation.

